# Genomic evidence of speciation by fusion linked to trophic niche expansion in a recent radiation of grasshoppers

**DOI:** 10.1101/2021.12.26.474180

**Authors:** Víctor Noguerales, Joaquín Ortego

**Author notes:** Corresponding author: Víctor Noguerales. - Instituto de Productos Naturales y Agrobiología (IPNA-CSIC), San Cristóbal de la Laguna, Tenerife, Islas Canarias, Spain.

## Abstract

Post-divergence gene flow can trigger a number of creative evolutionary outcomes, ranging from the transfer of beneficial alleles across species boundaries (*i.e*., adaptive introgression) to the formation of new species (*i.e*., hybrid speciation). While neutral and adaptive introgression has been broadly documented in nature, hybrid speciation is assumed to be rare and the evolutionary and ecological context facilitating this phenomenon still remains controversial. Through combining genomic and phenotypic data, we evaluate the hypothesis that the dual feeding regime (scrub legumes and gramineous herbs) of the taxonomically controversial grasshopper *Chorthippus saulcyi algoaldensis* resulted from hybridization between two sister taxa that exhibit contrasting host-plant specializations: *C. binotatus* (scrub legumes) and *C. saulcyi* (gramineous herbs). Genetic clustering analyses and inferences from coalescent-based demographic simulations confirmed that *C. s. algoaldensis* represents a uniquely evolving lineage and supported the ancient hybrid origin of this taxon (*ca*. 1.4 Ma), which provides a mechanistic explanation for its broader trophic niche and sheds light on its uncertain phylogenetic position. We propose a Pleistocene hybrid speciation model where range shifts resulting from climatic oscillations can promote the formation of hybrid swarms and facilitate its long-term persistence through geographic isolation from parental forms in topographically complex landscapes.

## INTRODUCTION

Gene flow is recognized as a fundamental control on the speciation process, with disparate evolutionary consequences that can impact either positively or negatively the formation and persistence of independently evolving lineages (Mallet 2007; Dynesius and Jansson 2014). At one extreme, gene flow can inhibit the onset of speciation by homogenizing gene pools (Slatkin 1985; Dynesius and Jansson 2014) and lead to ‘speciation reversal’ if lineages that have remained isolated for extended periods of time merge back into one after secondary contact (Seehausen et al. 2008; Kleindorfer et al. 2014; Kearns et al. 2018). At the opposite extreme, gene flow can generate a wide spectrum of creative evolutionary outcomes, ranging from adaptive introgression across species boundaries (Hedrick 2013; Suarez-Gonzalez et al. 2018a) to the formation of new hybrid species (Mallet 2007). Introgressive hybridization as a source of novel alleles conferring advantages to the recipient species has been widely documented in numerous organism groups (Hedrick 2013; Suarez-Gonzalez et al. 2018b). This phenomenon can lead to the acquisition of new traits, including the capacity to exploit new host-plants (Aardema and Andolfatto 2016), Müllerian mimicry (Pardo-Díaz et al. 2012; Enciso-Romero et al. 2017), and resistance to herbivores (Whitney et al. 2006), and has been proven to be instrumental in niche expansions (Scascitelli et al. 2010; Malinsky et al. 2018) and adaptation to suboptimal environmental conditions (Pfennig et al. 2016; Suarez-Gonzalez et al. 2018a; Leroy et al. 2020). In other cases, hybridization promotes the formation of new species (*i.e*., hybrid speciation) through the emergence of evolutionary innovations and reproductive isolation between parental and hybrid lineages (Gross and Rieseberg 2005; *e.g*., Gompert et al. 2006; Nice et al. 2013). Beyond a sporadic and fortuitous phenomenon, hybridization and introgression have been hypothesized to be responsible of fueling (“Syngameon” hypothesis; Seehausen 2004; Seehausen et al. 2014; *e.g*., Patton et al. 2020) or even igniting the onset of adaptive radiations (“Hybrid swarm origin of adaptive radiation” hypothesis; Meier et al. 2017).

Although introgression is rampant across the Tree of Life and allopolyploid hybrid speciation is relatively frequent in plants, homoploid hybrid speciation – speciation via hybridization without a change in chromosome number – has been much more rarely documented (Mallet 2007; Schumer et al. 2014; Taylor and Larson 2019). The major challenge for homoploid hybrid species to persist through evolutionary time is eluding the homogenizing effects of gene flow with their sympatric progenitors. Despite its evolutionary significance, the specific evolutionary and ecological contexts facilitating hybrid speciation are still controversial from both a theoretical and empirical perspective (Buerkle et al. 2000; Servedio et al. 2013; Schumer et al. 2014). What is well understood is that hybridization should have direct consequences on fitness of the incipient hybrid species through the emergence of novel or intermediate phenotypes on which natural or sexual selection can act on (Mavárez et al. 2006; Hedrick 2013; Taylor and Larson 2019). Literature on homoploid hybrid speciation has linked this phenomenon to rapid reproductive isolation between parental and hybrid lineages through strong assortative mating (Mavárez et al. 2006; Melo et al. 2009), colonization and adaptation to novel habitats with extreme conditions (Rieseberg et al. 1996, 2003; Gompert et al. 2006) or exploitation of new trophic resources (Schwarz et al. 2005; Lamichhaney et al. 2018;). However, successful homoploid hybrid speciation generally implies the co-occurrence of ecological circumstances (*e.g*., opening and colonization of a new niche space unavailable to either parental species) and genetic mechanisms (*e.g*., recombinant or transgressive phenotypes, chromosomal rearrangements, *etc*.) that increase the capacity to exploit novel resources and promote reproductive isolation (McCarthy et al. 1995; Gross and Rieseberg 2005; Schumer et al. 2014; Lamichhaney et al. 2018).

The species group *Chorthippus* (*Glyptobothrus*) (*binotatus*) (Charpentier, 1825) is a recently diverged complex of grasshoppers distributed in the westernmost portion of the Palearctic (Cigliano et al. 2017; Figs. 1-2). The complex is composed by eight taxa grouped in two major clades – *C. binotatus* and *C. saulcyi* clades – that exhibit distinct host-plant associations (Defaut 2011; Noguerales et al. 2018b; Fig. 1). While taxa from the clade *C. binotatus* exclusively feed on scrub legumes (Fabaceae, tribe Genisteae), lineages within the clade *C. saulcyi* show a feeding regime based on gramineous herbs (Poaceae; Picaud et al. 2003; Defaut 2011). The only exception is *C. saulcyi algoaldensis*, a narrow-endemic taxon distributed in the Massif Central (France; Fig. 2) that feeds indistinctly on scrub legumes and gramineous herbs (Defaut 2011; Noguerales et al. 2018b; Fig. 1). This taxon also presents a distinctive male calling song structure and an intermediate morphological position between its putative (*C. saulcyi*) and sister (*C. binotatus*) clades (Defaut 2011). While a preliminary study strongly supported the distinctiveness of each taxon within the complex, the phylogenetic placement of *C. s. algoaldensis* as a sister lineage to the rest of taxa within either *C. saulcyi* or *C. binotatus* clades remained unresolved (Noguerales et al. 2018b). Collectively, all these pieces of evidence raise the hypothesis of introgressive hybridization or speciation by fusion as the mechanistic explanation for the broader trophic niche of *C. s. algoaldensis* and its uncertain phylogenetic position, an evolutionary history departing from expectations under a strictly bifurcating model of divergence that should have left a distinctive signature on its genome (Meng and Kubatko 2009). Given that the rapid history of diversification of the complex is likely explained by processes of allopatric speciation during the Pleistocene (Mayer et al. 2010; Noguerales et al. 2018a, 2018b), distributional shifts driven by climatic fluctuations could have also provided ample opportunities for secondary contact and admixture among recently diverged lineages in which reproductive barriers to gene flow might be incomplete or absent (Hewitt 1999; Nolen et al. 2020; Ortego and Knowles 2021).

**Figure. 1.**
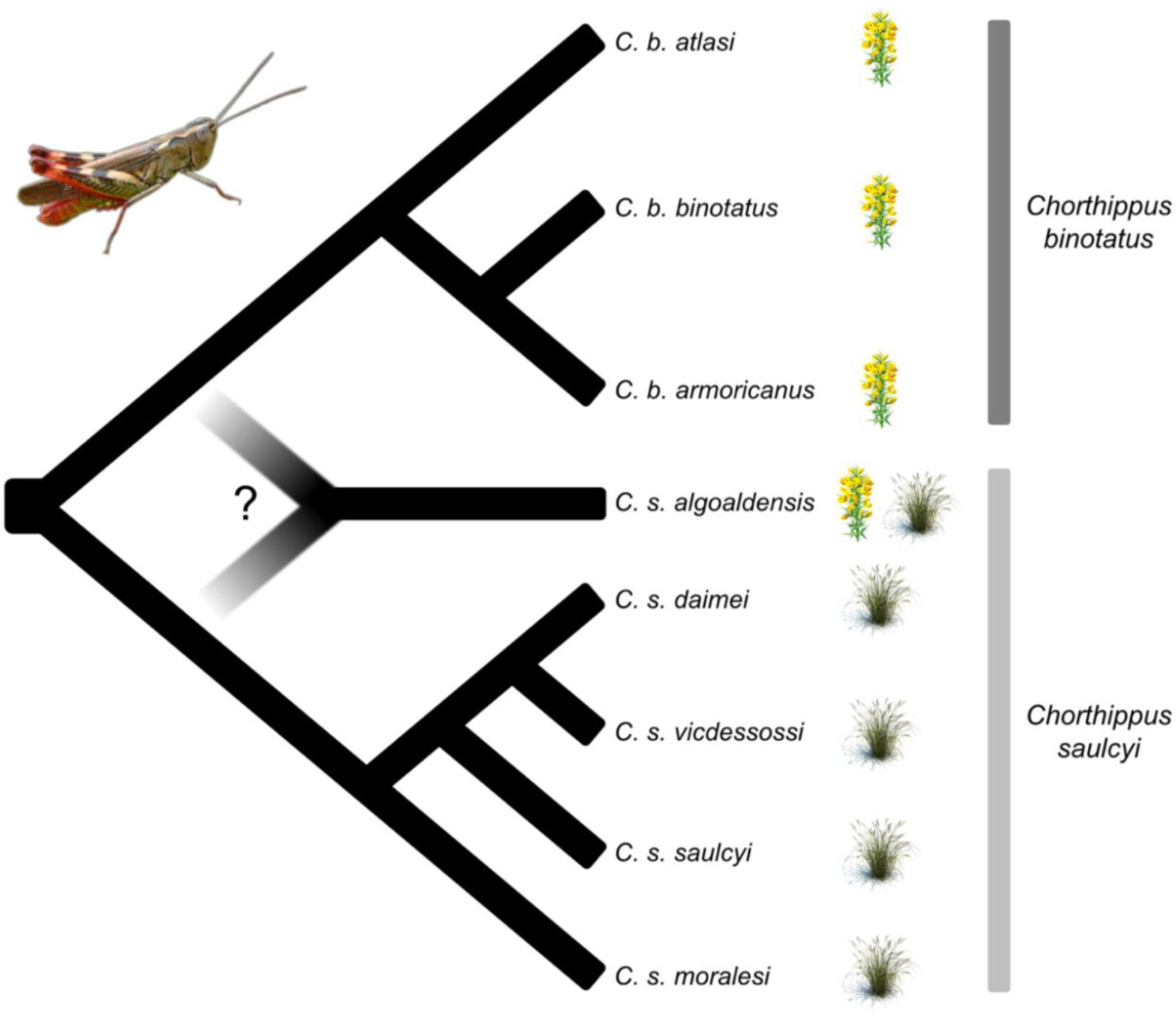
Schematic showing the phylogenetic relationships and host plant associations (scrub legumes *vs*. gramineous herbs) for the different taxa within the studied species complex and the taxonomic and phylogenetic uncertainties around the focal taxon *C. s. algoaldensis*. In this study we test alternative hypotheses concerning the roles of introgression and hybridization in the evolutionary history of *C. s. algoaldensis*, which has been traditionally included within *C. saulcyi* according to its morphology but shows dual host plant associations and presents an uncertain phylogenetic placement (Noguerales et al. 2018b).

**Figure 2.**
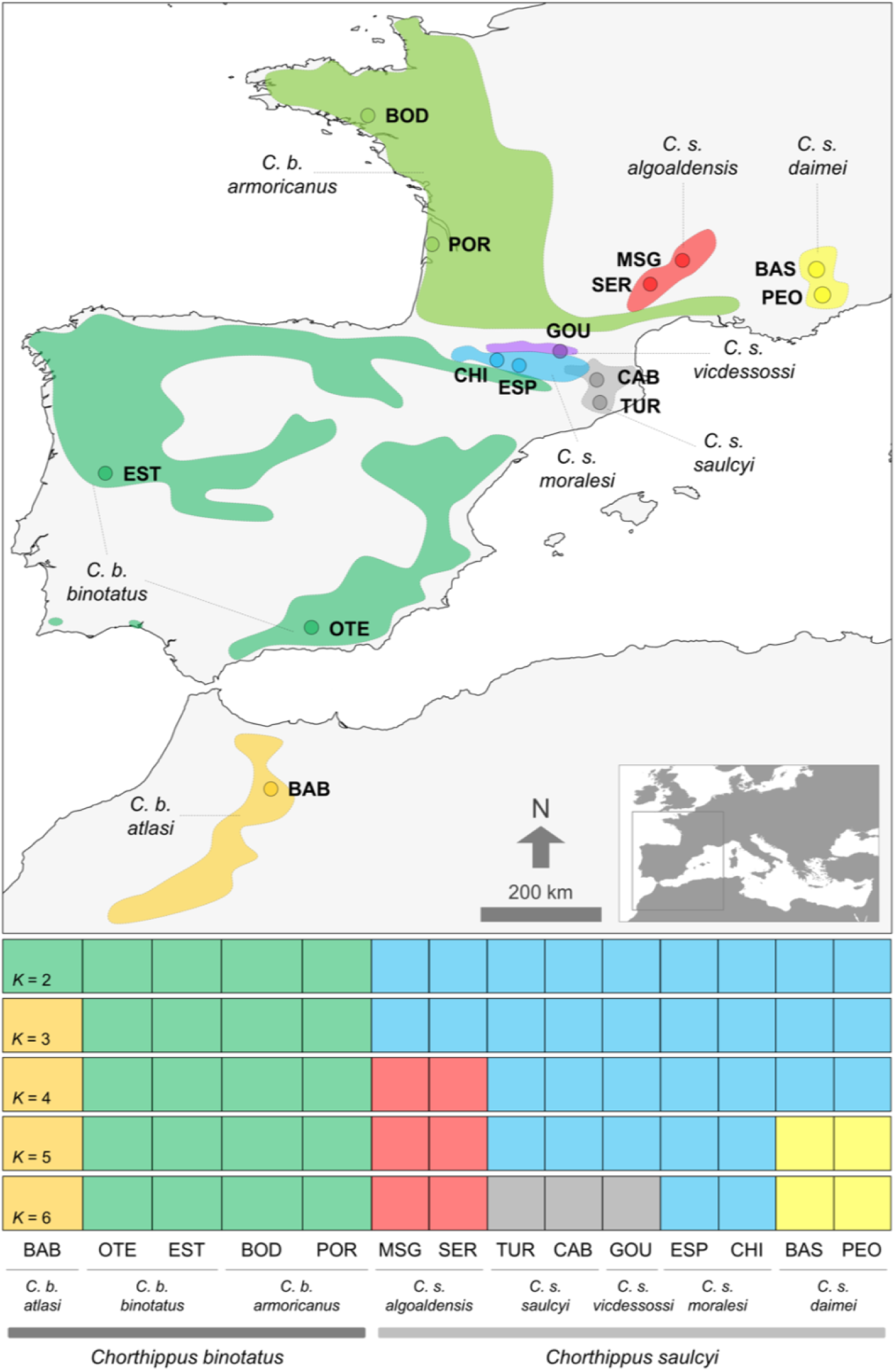
Geographical location of the sampled populations and distribution range of the different taxa from the studied species complex. Panels on the bottom show inferred genetic clustering from *K* = 2 to *K* = 6, the best-supported solutions as inferred by FASTSTRUCTURE. Individuals are partitioned into *K* colored segments representing the probability of belonging to the cluster with that color. Thin vertical black lines separate different populations. Population codes as in Table S1.

In this study, we integrate genomic data obtained through double-digest restriction associated DNA sequencing (ddRADseq) and phenotypic information to test the hypothesis that the dual feeding regime of *C. s. algoaldensis* represents a trophic niche expansion linked to hybridization between parental lineages with narrower host-plant requirements. Leveraging inferences obtained through both phylogenomics and population genetics frameworks, we evaluate the specific pathways that might have led to the evolution of the contrasting feeding strategies within the studied species complex. First, we perform clustering analyses to quantify patterns of population genetic structure, detect signatures of admixture across species boundaries, and determine the genetic distinctiveness and cohesiveness of putative taxa within the complex. Second, we test whether the unresolved phylogenetic position of *C. s. algoaldensis* is explained by incomplete lineage sorting or introgression and congruent with phenotypic variation. Finally, we use a model-based simulation approach to test refined hypotheses invoking alternative scenarios of speciation, namely strict bifurcation, introgression, and hybrid speciation, and infer the mode and estimate the timing of species formation.

## MATERIALS AND METHODS

### Sample collection

We collected samples from the eight putative taxa (14 populations, 231 individuals) constituting the species group *Chorthippus* (*Glyptobothrus*) *binotatus* (Charpentier, 1825) (Noguerales et al. 2018b and references therein; Table S1; Fig. 2). When possible, we collected two populations per taxon and tried to maximize the distance from each other in order to include samples representative of their respective distribution ranges (Table S1; Fig. 2).

### Genomic data

We processed extracted DNA from a subset of 77 individuals (5-7 individuals per population; Table S1) into two genomic libraries following the double digestion restriction associated DNA sequencing (ddRADseq) procedure described in Peterson et al. (2012). We also included into the libraries six individuals from *Chorthippus* (*Glyptobothrus*) *biroi* (Kuthy, 1907) (Table S1), which were used as an outgroup in phylogenomic and introgression/hybridization analyses. Details on the preparation of ddRADseq libraries are presented in Methods S1 (Supporting Information). Raw sequences were demultiplexed and pre-processed using STACKS v.1.35 (Catchen et al. 2013) and assembled in PYRAD v.3.0.66 (Eaton 2014). Methods S2 in Supporting Information provides all details on data filtering and sequence assembling.

### Genetic clustering analyses

Genetic clustering of the studied taxa and populations was inferred using the variational Bayesian framework implemented in FASTSTRUCTURE v.1.0 (Raj et al. 2014). Ten independent replicates were performed for a range of different *K* genetic clusters (*K* = 1-14) using the simple prior and a convergence criterion of 1 × 10^-7^. Following Raj et al. (2014), the number of genetic clusters that best describes our data was assessed by calculating the metrics *K*\*_ø_^c^, the value of *K* that maximizes log-marginal likelihood lower bound of the data, and *K*\*_ε_, the smallest number of model components explaining at least 99% of cumulative ancestry contribution in our sample.

### Phylogenomic inference

Phylogenomic relationships among taxa were reconstructed using two different coalescent-based methods for species tree estimation. First, we ran SVDQUARTETS (Chifman and Kubatko 2014) including *C. biroi* as an outgroup, exhaustively evaluating all possible quartets, and performing nonparametric bootstrapping with 100 replicates for quantifying uncertainty in relationships. Second, we used the Bayesian coalescent model implemented in SNAPP v.1.3 (Bryant et al. 2012). We applied two alternative gamma distributions for the ancestral population size parameter (θ), namely G(2, 200) and G(2, 2000), and leaving default settings for all other parameters. Due to high computational burden of SNAPP analyses, the number of taxa partitions was limited by including only one population per taxon. We used TRACER v.1.4 to examine log files and check stationarity and convergence of the chains and confirm that effective sample sizes (ESS) for all parameters were >200. We removed 10% of trees as burn-in and combined tree and log files for replicated runs using LOGCOMBINER v.2.4.7. Maximum credibility trees were obtained using TREEANNOTATOR v.2.4.7 and the full set of likely species trees was displayed with DENSITREE v.2.2.6, which is expected to show fuzziness in parts of the tree due to gene flow or other causes of phylogenetic conflict (Bouckaert 2010).

### Testing for introgression

We used four-taxon ABBA/BABA tests based on the *D*-statistic to determine the role of hybridization/introgression on explaining unresolved phylogenetic relationships involving *C. s. algoaldensis* (Durand et al. 2011). This method enables evaluating to what extent gene-tree incongruences have resulted from either gene flow between non-sister taxa or retention of ancestral genetic variation (*i.e*., incomplete lineage sorting, ILS). Assuming that the sister species P_1_ and P_2_ diverged from P_3_ and an outgroup species O, the *D*-statistic is used to test the null hypothesis of no introgression (*D* = 0) between P_3_ and P_1_ or P_2_. *D*-values significantly different from zero indicate gene flow between P_1_ and P_3_ (*D* <0) or between P_2_ and P_3_ (*D* >0). We assigned taxa within *C*. *saulcyi* (excluding *C. s. algoaldensis*) to P_1_, *C. s. algoaldensis* to P_2_, taxa within *C*. *binotatus* to P_3_, and *C. biroi* to the outgroup (O). We performed ABBA/BABA tests in PYRAD and used 1000 bootstrap replicates to obtain the standard deviation of the *D*-statistic (Eaton and Ree 2013). We ran ABBA/BABA tests combining data from all taxa and populations in each group (P_1_, P_2_ and P_3_) and also performing independent analyses considering all possible taxa combinations (*i.e.*, subspecies within *C. binotatus* and *C. saulcyi*; Table S2).

We complemented ABBA/BABA tests with analyses based on the recently developed software HYDE (Blischak et al. 2018), which tests for hybridization using phylogenetic invariants under a model of hybrid speciation accommodating coalescent stochasticity (Kubatko and Chifman 2019). This approach estimates the amount of admixture (γ) in a putative hybrid population (P_HYB_), which is modelled as a mixture of two parental populations, P_1_ (1-γ) and P_3_ (γ). Conversely to other similar approaches, HYDE performs individual-based hypothesis tests and bootstrap resampling within the putative hybrid population (Blischak et al. 2018). Consequently, this method allows detecting heterogeneous admixture patterns across individuals within the focal population and detecting ongoing hybridization processes. We ran HYDE assuming the same four-taxon topology used for ABBA/BABA tests, in this case assigning taxa within *C. saulcyi* (excluding *C. s. algoaldensis*) to P_1_, taxa within *C. binotatus* to P_3_, *C. s. algoaldensis* to the putative hybrid lineage (P_HYB_), and *C. biroi* to the outgroup. Estimates of γ parameter were calculated across 100 bootstrap resampling replicates of the individuals from the hybrid group using the *bootstrap_hyde.py* script. Heterogeneity in the degree of admixture across putatively hybrid individuals (γ_IND_) was estimated using the *individual_hyde.py* script, which runs separate analyses on each individual belonging to P_HYB_ (individual-based analyses). We ran HYDE using the same taxa combinations as for ABBA/BABA tests (Table S3).

### Testing alternative demographic models

We evaluated alternative models of speciation for *C. s. algoaldensis* in order to determine whether its uncertain phylogenetic position when assuming a strictly bifurcating tree is a consequence of an introgression event from *C. binotatus* into *C. s. algoaldensis* (*i.e*., a pulse of gene flow) after the latter diverged from *C. saulcyi* (hereafter, “introgression model”) or if, alternatively, *C. s. algoaldensis* originated from an admixture event between *C. binotatus* and *C. saulcyi* (hereafter, “speciation by fusion model”; *sensu* Grant and Grant 2018; see also Barrera-Guzmán et al. 2018). These two alternative models, together with a null model considering a strictly bifurcating history of divergence (hereafter, “strictly bifurcating model”), were built both assuming no migration among demes and considering post-divergence gene flow (*i.e*., a total of six models, illustrated in Fig. 4). To evaluate the relative statistical support for each of these alternative demographic scenarios, we estimated the composite likelihood of the observed data given a specified model using the site frequency spectrum (SFS) and the simulation-based approach implemented in FASTSIMCOAL2 v.2.5.2.21 (Excoffier et al. 2013). All sampling populations were included into the analyses and assigned to one of the three demes considered (*C. binotatus*, *C. saulcyi*, and the putative hybrid taxon *C. s. algoaldensis*) according to phylogenomic inferences (see Results section).

**Figure 4.**
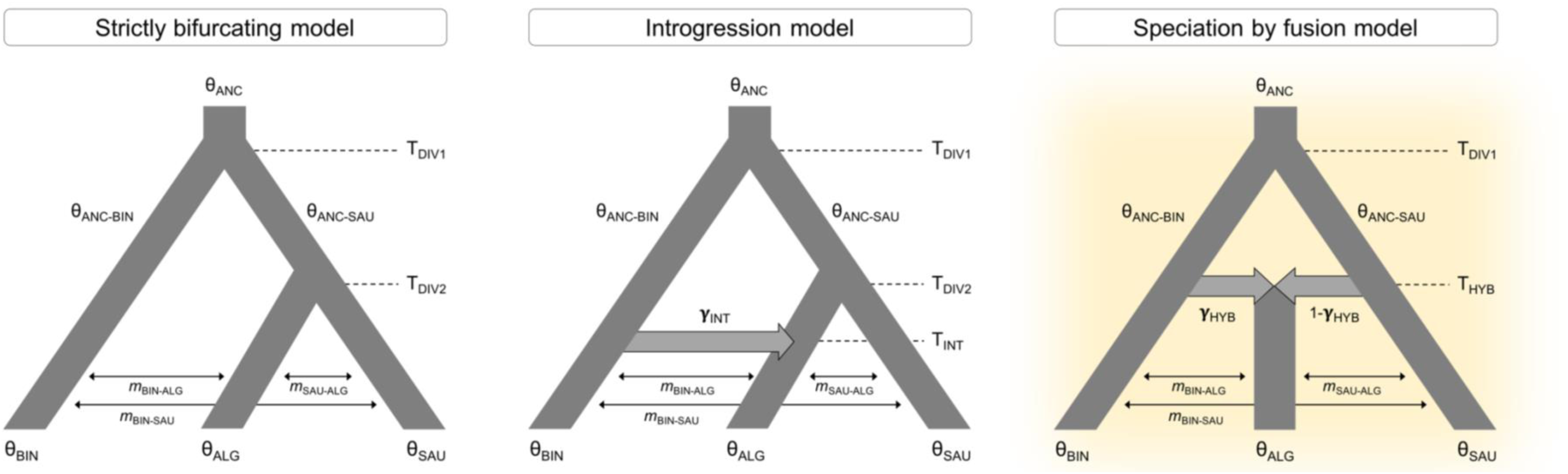
Alternative demographic scenarios tested using FASTSIMCOAL2, including strictly bifurcating, introgression, and speciation by fusion models. Models were tested both considering and not considering post-divergence or post-hybridization gene flow (see Table 1). Model parameters include ancestral (θ_ANC_, θ_ANC-SAU_, θ_ANC-BIN_) and contemporary (θ_SAU_, θ_ALG_, θ_BIN_) effective population sizes, timing of divergence (T_DIV1_, T_DIV2_), introgression (T_INT_) and hybridization (T_HYB_), introgression (γ_INT_) and hybridization (γ_HYB_) coefficients, and migration rates per generation (*m*). The best-supported model is highlighted.

**Table 1.** Comparison of alternative models (see Fig. 4) tested using FASTSIMCOAL2. The three main models were built both considering and not considering post-divergence or post-hybridization gene flow among demes. The best-supported model is highlighted in bold.

| Model | Gene flow | lnL | *k* | AIC | ΔAIC | $\omega_i$ |
| --- | --- | --- | --- | --- | --- | --- |
| Strictly bifurcating | No | -3158.59 | 6 | 6329.19 | 121.19 | 0.00 |
| Introgression | No | -3147.64 | 8 | 6312.31 | 103.62 | 0.00 |
| Speciation by fusion | No | -3148.15 | 8 | 6313.84 | 104.65 | 0.00 |
| Strictly bifurcating | Yes | -3101.70 | 9 | 6221.39 | 13.74 | 0.00 |
| Introgression | Yes | -3097.91 | 11 | 6217.82 | 10.17 | 0.01 |
| **Speciation by fusion** | **Yes** | **-3092.83** | **11** | **6207.65** | **0.00** | **0.99** |

lnL, maximum likelihood estimate of the model; *k*, number of parameters in the model; AIC, Akaike's information criterion value; ΔAIC, difference in AIC value from that of the strongest model; $\omega_i$ , AIC weight.

We calculated a folded joint SFS using the *easySFS.py* script (I. Overcast, https://github.com/isaacovercast/easySFS). We considered a single SNP per locus to avoid the effects of linkage disequilibrium and down sampled each population group (deme) to 50% of individuals to remove all missing data for the calculation of the joint SFS, minimize errors with allele frequency estimates, and maximize the number of variable SNPs retained. The final SFS contained 1998 variable SNPs. Because we did not include invariable sites in the SFS, we used the “removeZeroSFS” option in FASTSIMCOAL2 and fixed the effective population size for one of the demes (*C. binotatus*) to enable the estimation of other parameters in FASTSIMCOAL2 (Excoffier et al. 2013; Papadopoulou and Knowles, 2015). The effective population size fixed in the model was calculated from the level of nucleotide diversity (π) and estimates of mutation rate per site per generation (μ), since *N*_e_ = (π/4μ). Nucleotide diversity (π) was estimated from polymorphic and non-polymorphic loci using DNASP v.6.12.03 (Rozas et al. 2017). We considered the mutation rate per site per generation of 2.8 × 10^-9^ estimated for *Drosophila melanogaster* (Keightley et al. 2014).

Each model was run 100 replicated times considering 100,000-250,000 simulations for the calculation of the composite likelihood, 10-40 expectation-conditional maximization (ECM) cycles, and a stopping criterion of 0.001 (Excoffier et al. 2013). We used an information-theoretic model selection approach based on the Akaike’s information criterion (AIC) to determine the probability of each model given the observed data (Burnham and Anderson 2002; *e.g*., Thomé and Carsterns 2016). After the maximum likelihood was estimated for each model in every replicate, we calculated the AIC scores as detailed in Thomé and Carsterns (2016). AIC values for each model were rescaled (ΔAIC) calculating the difference between the AIC value of each model and the minimum AIC obtained among all competing models (*i.e*., the best model has ΔAIC = 0). Point estimates of the different demographic parameters for the best supported model were selected from the run with the highest maximum composite likelihood. Finally, we calculated confidence intervals (based on the percentile method; *e.g*., de Manuel et al. 2016) of parameter estimates from 100 parametric bootstrap replicates by simulating SFS from the maximum composite likelihood estimates and re-estimating parameters each time (Excoffier et al. 2013).

### Phenotypic variation analyses

In order to assess the effect of hypothetical genetic admixture between parental taxa on the phenotype of the putative hybrid lineage (*C. s. algoaldensis*), we took digital images of taxonomically relevant traits (left hind femur, left forewing and pronotum; Defaut 2011; Noguerales et al. 2018b) from each individual included in the study (Table S1) and analyzed them by means of both linear and geometric morphometric approaches. For the linear morphology approach, we focused on three ratio traits that have been already considered in previous taxonomic studies of the group (Defaut 2011; Noguerales et al. 2018b), namely: (i) forewing length relative to femur length (FWL/FL), (ii) forewing median area length relative to total forewing length (MAL/FWL), and (iii) prozone length relative to total pronotum length (PZ/PR). Regarding the geometric morphometric approach, we focused on forewing shape (*e.g.*, Klingenberg et al. 2010; Noguerales et al. 2018b; Tonzo et al. 2019). Variation in forewing shape was analyzed in MORPHOJ v.1.05d (Klingenberg 2011) considering ten homologous landmarks that have been previously shown to be highly informative in describing geometric morphometric variation of this trait within the study group (Noguerales et al. 2016, 2018b). Briefly, we conducted a Procrustes fit separately for each sex, removed the allometry effect on trait shape, and summarized size-corrected shape variation by means of principal component analyses (PCA) (for more details, see Klingenberg 2011). Differences among subspecies and among the three main groups (*C. s. algoaldensis* and its putative parental taxa *C. binotatus* and *C. saulcyi*) for ratio traits were tested using one-way ANOVAs in R v.4.0.3 (R Core Team 2021). Likewise, differences among subspecies and the three main groups in forewing shape was assessed by calculating Mahalanobis distances (*D*) from a canonical variate analyses (CVA) and conducting 10,000 permutation tests to calculate statistical significance (Klingenberg 2011). All traits were analyzed separately for each sex due to the considerable sexual size dimorphism in Orthoptera (Hochkirch and Gröning 2008).

## RESULTS

### Genomic data

Illumina sequencing provided a total of 211.01 M sequences reads, with an average 2.45 M sequence reads per individual (SD = 0.39 M) (Fig. S1). After the different filtering and assembly steps, each individual retained on average 2.15 M sequence reads (SD = 0.35 M) (Fig. S1). Finally, the within-sample clustering step yielded on average 56,777 loci (SD = 7707) per individual.

### Genetic clustering analyses

Genetic clustering analyses in FASTSTRUCTURE supported *K* = 2 and *K* = 6 as the most likely number of genetic groups according to the metrics *K*\*_ø_^c^ and *K*\*_ε_, respectively. When considering *K* = 2, the different taxa were assigned to either *C. binotatus* or *C. saulcyi* in concordance with the prevailing taxonomy of the group. For *K* = 6, *C. saulcyi* split into four genetic clusters that corresponded to *C. s. algoaldensis, C. s. moralesi, C. s. daimei*, and *C. s. saulcyi* together with the subspecies *C. s. vicdessossi*, whereas *C. binotatus* divided into two well-defined genetic clusters corresponding to the Maghrebian (*C. b. atlasi*) and European (*C. b. binotatus* and *C. b. armoricanus*) taxa (Fig. 2). Evaluation of alternative *K*-values within the range of best-supported clustering solutions (*K* = 2-6) confirmed that genomic variation is hierarchically organized and infraspecific entities (*i.e*., subspecies and populations) are nested into well-defined genetic clusters corresponding to the different putative taxa. None clustering solution showed evidence of genetic admixture at species, subspecies or population levels (ancestry >99.99%; Fig. 2). Inferences assuming an increasing number of genetic clusters (*K* >6) showed no further genetic structure and consistently yielded “ghost clusters” (*i.e*., clusters with no population or individual assigned to them; Guillot et al. 2005).

### Phylogenomic inference

The species tree reconstructed in SVDQUARTETS revealed the existence of two major clades corresponding to the *C. binotatus* and *C. saulcyi* groups. In line with inferences from genetic clustering analyses, populations from the same putative subspecies grouped into well-supported monophyletic sub-clades (Fig. 3). However, whereas the relationships among subspecies from the *C. binotatus* group were well resolved (node support >99%), the phylogenetic position of *C. s. algoaldensis* was unclear and the node separating it from the rest of taxa within the *C. saulcyi* group was the one showing the lowest bootstrapping support (= 79 %) in the whole tree (Fig. 3). The topology inferred by SNAPP was similar to that from SVDQUARTETS and also showed that the phylogenetic relationships within the *C. saulcyi* group were not well supported (Fig. 3; Fig. S2). However, in this case the split concerning the focal taxon *C. s. algoaldensis* showed no evidence of topological uncertainty (Fig. S2). SNAPP runs considering different priors for the θ parameter converged on the same topology.

**Figure 3.**
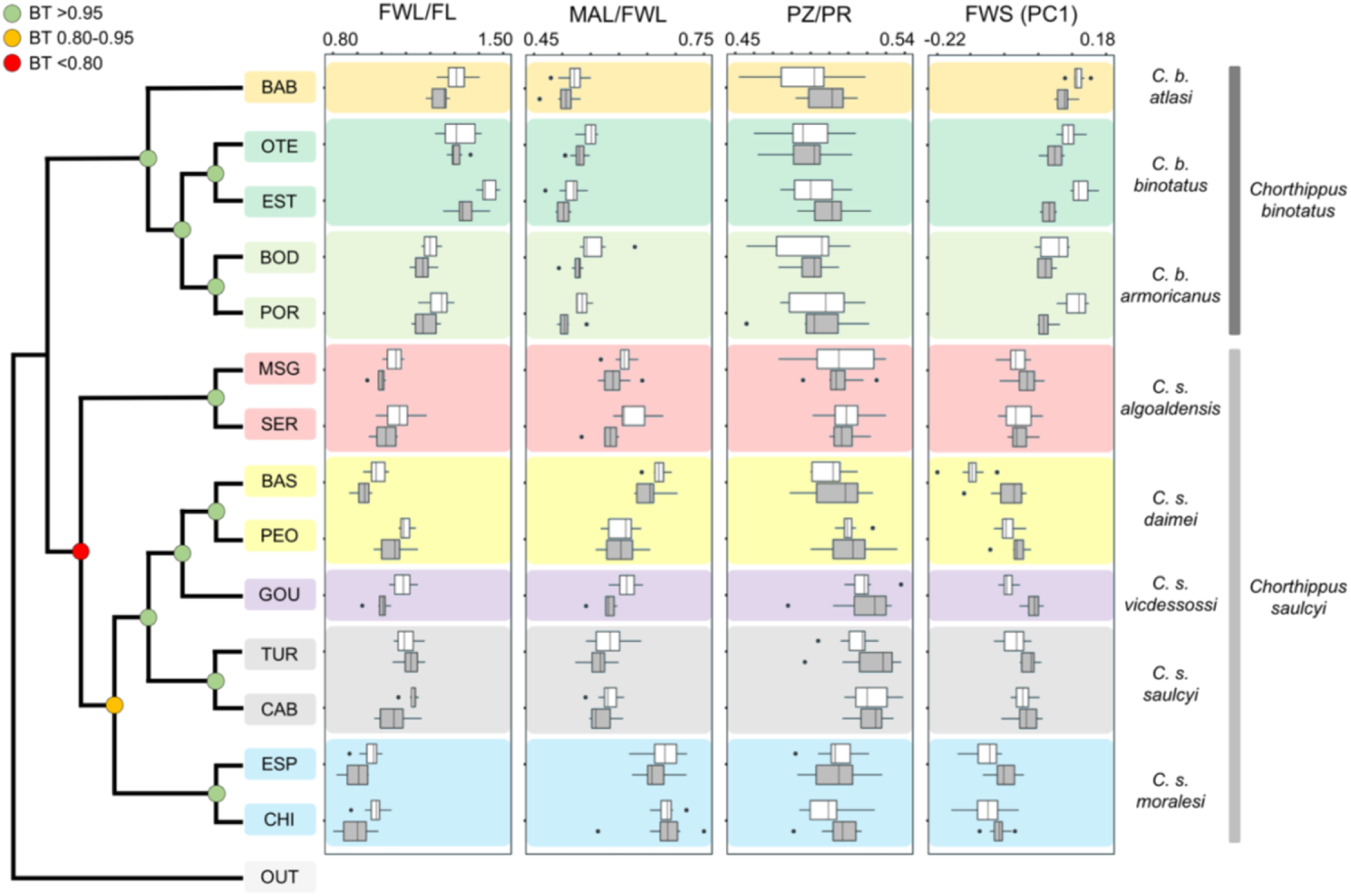
Species tree inferred with SVDQUARTETS showing the phylogenetic relationships among the different populations and taxa from the studied species complex. Bootstrapping values (BT) are indicated on the nodes using different colors as detailed in the legend. Boxplots on the right panels summarize phenotypic variation for each studied trait, including forewing length relative to femur length (FWL/FL), forewing median area length relative to forewing length (MAL/FWL), prozone length relative to pronotum length (PZ/PR), and forewing shape (FWS) variation based on the first principal component (PC1). Outliers are shown as black dots and white and grey boxplots represent males and females, respectively. Population codes as in Table S1.

### Testing for introgression

Results of *D*-statistic tests revealed significant introgression involving *C. s. algoaldensis* and *C. binotatus* (*D*_s_ = 0.43 ± 0.06 SD; BABA = 65; ABBA = 103; *Z*-score = 7.56; *P*-value <0.001; number of loci = 1245). Similar results were obtained when analyses were performed considering all possible taxa combinations (*i.e*., subspecies within *C. binotatus* and *C. saulcyi*; Table S2). In concordance with *D*-statistic tests, HYDE analyses detected high levels of admixture in *C. s. algoaldensis* (γ = 0.48; 95% CIs = 0.47-0.49; *Z*-score = 55.24; *P*-value <0.001). Analyses performed considering all possible taxa combinations provided similar inferences (Table S3). Individual-based analyses were all highly significant and yielded similar estimates of γ parameter across individuals and populations within *C. s. algoaldensis* (γ_IND_ range = 0.46-0.50, all *Z*-score >31.34; all *P*-value <0.001), indicating a uniform level of genetic admixture likely resulted from an ancient hybridization event (Blischak et al. 2018). Similar γ_IND_ estimates were obtained when analyses were performed considering all possible taxa combinations (Table S3).

### Testing alternative demographic models

FASTSIMCOAL2 analyses identified the speciation by fusion model incorporating post-divergence gene flow as the most likely scenario (Table 1; Fig. 4), supporting that *C. s. algoaldensis* originated from a hybridization event between *C. binotatus* and *C. saulcyi*. The introgression model incorporating post-divergence gene flow was the second-most supported scenario, although it cannot be considered statistically equivalent to the most likely scenario (ΔAIC = 10.17; Table 1). The strictly bifurcating model (ΔAIC >13) and models that did not incorporate post-divergence gene flow were highly unlikely (ΔAIC >100; Table 1). Considering that the studied taxa are univoltine (*i.e.*, 1-year generation time), FASTSIMCOAL2 estimated that *C. binotatus* and *C. saulcyi* diverged from a common ancestor (T_DIV1_) *ca*. 2.1 Ma (95% CI: 1.3-2.2 Ma; Table 2). The hybridization event (T_HYB_) that led to the formation of *C. s. algoaldensis* was estimated to have occurred *ca*. 1.4 Ma (95% CI: 1.0-1.6 Ma; Table 2). These analyses showed that *ca*. 24% of alleles (95% CI: 10-43%) of *C. s. algoaldensis* were inherited (γ_HYB_) from *C. binotatus* and post-hybridization migration rates per generation (*m*) among all demes were consistently very low (Table 2).

**Table 2.** Parameters inferred from coalescent simulations with FASTSIMCOAL2 under the best-supported speciation by fusion model (see Fig. 4). For each parameter, we show its point estimate and lower and upper 95% confidence intervals. Model parameters include ancestral (θ_ANC_, θ_ANC-SAU_, θ_ANC-BIN_) and contemporary (θ_SAU_, θ_ALG_) mutation-scaled effective population sizes, timing of ancestral divergence (T_DIV1_) and hybridization (T_HYB_), admixture coefficient (γ_HYB_) and migration rates per generation (m) among demes. Note that the effective population size of C. binotatus (θ_BIN_) is not presented because it was fixed in FASTSIMCOAL2 analyses to enable the estimation of other parameters.

| Parameter | Point estimate | Lower bound | Upper bound |
| --- | --- | --- | --- |
| $\theta_{\text{ANC}}$ | 40,956 | 33,427 | 527,869 |
| $\theta_{\text{ANC-SAU}}$ | 933,380 | 308,449 | 1,617,713 |
| $\theta_{\text{ANC-BIN}}$ | 9721 | 4915 | 272,882 |
| $\theta_{\text{ALG}}$ | 680,545 | 547,955 | 738,974 |
| $\theta_{\text{SAU}}$ | 2,834,490 | 2,329,980 | 3,079,558 |
| $T_{\text{HYB}}$ | 1,365,495 | 1,019,168 | 1,567,835 |
| $T_{\text{DIV1}}$ | 2,052,400 | 1,348,179 | 2,195,238 |
| $\gamma_{\text{HYB}}$ | 0.24 | 0.10 | 0.43 |
| $m_{\text{SAU-ALG}}$ | $7.19 \times 10^{-8}$ | $4.45 \times 10^{-8}$ | $1.06 \times 10^{-7}$ |
| $m_{\text{BIN-ALG}}$ | $2.74 \times 10^{-8}$ | $1.18 \times 10^{-8}$ | $4.05 \times 10^{-8}$ |
| $m_{\text{BIN-SAU}}$ | $5.74 \times 10^{-8}$ | $4.08 \times 10^{-8}$ | $7.09 \times 10^{-8}$ |

### Phenotypic variation analyses

Regarding linear morphological analyses, one-way ANOVAs showed significant differences among the main groups (*C. binotatus*, *C. saulcyi* and *C. s. algoaldensis*) and subspecies for the three ratio traits in both sexes (all *P*-values <0.001). Post-hoc Tukeýs tests at the group level revealed that only comparisons between *C. binotatus* and any of the other two groups (*C. saulcyi* and *C. s. algoaldensis*) showed significant differences (*P*-values <0.05) for any trait and sex (Table S4; Fig. 3). Post-hoc tests at subspecies level showed that *C. s. algoaldensis* was significantly different from any taxon within *C. binotatus* for the forewing-derived ratio traits (FWL/FL and MAL/FWL) in both sexes (all *P*-values <0.05; Table S5; Fig. 3). Within *C. saulcyi,* we also found significant differences among taxa for these two ratio traits in both sexes, particularly in comparisons involving *C. s. algoaldensis* and those subspecies that are specially short-winged (*C. s. daimei* and *C. s. moralesi*) (Table S5; Fig. 3). The pronotum-derived ratio trait (PZ/PR) only showed significant differences in post-hoc tests for a few comparisons, mainly when subspecies *C. b. atlasi and C. s. saulcyi* were involved (Table S5).

Morphometric geometric analyses on forewing shape showed that *C. s. algoaldensis* clustered within the rest of the *C. saulcyi* group and that the overlapping between *C. binotatus* and *C. saulcyi* groups was low (Fig. 3; Fig. S3). Although Mahalanobis distances (*D*) were significantly different between all groups and subspecies for both sexes (Tables S6-S7), greater differences were found between *C. binotatus* and *C. s. algoaldensis* than when this taxon was compared to *C. saulcyi* (Tables S6-S7).

## DISCUSSION

We present evidence for hybridization to be a key component in the diversification of a species complex of Gomphocerinae grasshoppers. Phylogenetic tests and inferences from coalescent-based demographic simulations supported the hybrid origin of the narrowly distributed taxon *C. s. algoaldensis*, shedding light on its uncertain taxonomic position and providing a mechanistic explanation for its dual host-plant feeding regime. Although we cannot categorically conclude that isolating mechanisms (*i.e*., speciation itself) were triggered by hybridization, our study offers clues for alternative scenarios of hybrid speciation complementing more conservative definitions of this phenomenon proposed in previous literature (McCarthy et al. 1995; Buerkle et al. 2000; Gross and Rieseberg 2005). Below, we discuss the underlying biogeographic scenario and ecological factors that may have promoted the formation of *C. s. algoaldensis* and its stabilization as an independently evolving lineage of hybrid origin.

### Hybrid speciation in *Chorthippus* grasshoppers?

It has been estimated that the genome of at least 10% of species of animals has been sculpted by episodes of interspecific gene flow (Mallet 2005, 2007). Our study adds to the accumulating evidence on this phenomenon by demonstrating that the grasshopper *C. s. algoaldensis* represents a relatively ancient lineage of hybrid origin. In line with inferences from phylogenetic tests (Tables S2-S3), coalescent-based demographic analyses strongly supported a speciation-by-fusion model (Table 1). This confirms the role of genetic admixture in shaping the evolutionary history *C. s. algoaldensis* while simultaneously offering an explanation for its uncertain phylogenetic placement and conflicting gene tree topologies in the complex (Fig. 2; Noguerales et al. 2018b). The fact that the genomic signatures of hybridization were revealed by phylogenetic tests involving each extant parental lineage (*i.e*., the different subspecies from each parental taxa; Tables S2-S3), indicates that the fusion event leading to the formation of *C. s. algoaldensis* was ancient and predated the diversification of *C. binotatus* and *C. saulcyi* into their respective infraspecific lineages (*e.g*., Ortego and Knowles 2021). In contrast to previous studies documenting gene flow across species boundaries in other grasshoppers (*e.g.*, Orr et al. 1994; Bridle et al. 2002; Nolen et al. 2020; Tonzo et al. 2020; Ortego and Knowles 2021), the recombinant genome of *C. s. algoaldensis* is compatible with hybrid speciation rather than with a scenario of bifurcating divergence followed by a pulse of gene flow (*i.e*., introgressive hybridization; Table 1). Thus, we argue that this taxon represents a putative case of homoploid hybrid speciation, as proposed for a handful of organisms including butterflies (Gompert et al. 2006), fruitflies (Schwarz et al. 2005), birds (Barrera-Guzman et al. 2018) and fishes (Keller et al. 2013).

In the last decade, the growing literature documenting presumably homoploid hybrid species has been accompanied by research efforts for outlining the criteria that should be satisfied to support this speciation mode, namely: (i) genomes exhibit signatures of hybridization (*criterion 1*), (ii) hybrids are reproductively isolated from parental forms (*criterion 2*), and (iii) isolating mechanisms were triggered by hybridization (*criterion 3*) (Abott et al. 2013; Schumer et al. 2014; but see Nieto-Feliner et al. 2017). Our genomic data clearly demonstrate that *C. s. algoaldensis* has a hybrid origin (*criterion 1*) and several lines of evidence suggest that contemporary populations of *C. s. algoaldensis* are reproductively isolated from the rest of taxa of the group (*criterion 2*). Even though its adjacent distribution with other taxa within the complex (*e.g*., *C. s. daimei* and *C. b. armoricanus*; with nearest populations <130 km; Fig. 2; Noguerales et al. 2018b) and, thus, likely opportunities for secondary contact resulting from range expansions promoted by the marked climatic oscillations that followed the speciation-by-fusion event (*ca*. 1.4 Ma; Table 2), we found no evidence for post-divergence gene flow between *C. s. algoaldensis* and parental taxa. In this line, clustering analyses revealed no signatures of recent genetic admixture among lineages (individual-based cluster memberships >0.999%; Fig. 2), inheritance probabilities from each progenitor varied little across individuals and populations (γ parameter; Table S3; *i.e*., populations are at genotypic equilibrium), and migration rates per generation between *C. s. algoaldensis* and parental taxa were extremely low (<8 × 10^-8^; Table 2). This supports the genetic cohesiveness and distinctiveness of *C. s. algoaldensis* and its long-term persistence as an independently evolving lineage (Barton and Hewitt 1985; *e.g*., Gompert et al. 2006, 2014).

Despite different lines of evidence satisfy or lend support to the two first criteria for homoploid hybrid speciation, we cannot determine whether reproductive isolation evolved as a direct consequence of hybridization (*criterion 3*; Schumer et al. 2014). Although this criterion is nearly impossible to demonstrate in ancient events leading to hybrid speciation, we hypothesize a scenario where coupled shifts in display traits and mating preferences resulted from hybridization could have played a relevant role on the reproductive isolation between *C. s. algoaldensis* and parental taxa (Rosentahl 2013). Gomphocerinae grasshoppers use acoustic signals for species recognition and mate choice (Nattier et al. 2011; Song et al. 2020), a behavior that has been extensively documented to be involved in reproductive isolation (Perdeck 1958; Bridle and Butlin 2002). Males of *C. s. algoaldensis* exhibit a slightly distinct courtship song relative to the rest of taxa within the group (Defaut 2011), raising the question of whether these differences emerged directly through hybridization and were instrumental as a prezygotic isolating barrier during the early stages of the nascent hybrid species. While it is also possible that song differences in *C. s. algoaldensis* evolved through mechanisms unrelated to hybridization (*e.g*., as a by-product of genetic drift due to long-term geographical isolation), the fact that the remaining taxa of the group produce very similar calling songs suggests that this alternative hypothesis is less plausible (Defaut 2011). The distinct courtship behavior of *C. s. algoaldensis* would be also in concordance with field and experimental studies revealing that male hybrids of Gomphocerinae grasshoppers exhibit novel, intermediate, or even more elaborated calling songs on which sexual section can act on (Perdeck 1958; Vedenina and von Helversen 2003; reviewed in Mayer et al. 2010). As suggested in other putative hybrid species (*e.g*., Naisbit et al. 2001; Mavárez et al. 2006; Melo et al. 2009; Lamichhaney et al. 2018), assortative mating could have favored the rapid establishment of reproductive barriers with parental taxa and provided the behavioral context for the hybrid lineage to progress toward speciation (Rosenthal 2013; Lamichhaney et al. 2018; see also Vedenina and von Helversen 2003).

### Phenotypic outcomes of hybridization

Demographic modelling estimated that the genome of *C. s. algoaldensis* presents a high level of admixed ancestry, which provides a mechanistic explanation for the phenotypic and ecological attributes of this taxon. The asymmetric genetic contribution from each parental lineage (*ca*. 24% and 76% of gene copies originated from *C. binotatus* and *C. saulcyi,* respectively; Table S1) is congruent with the fact *C. s. algoaldensis* tends to exhibit a greater morphological affinity with taxa belonging to its putative species (*C. saulcyi*) than with lineages of *C. binotatus* (Figs. 2-3; Fig. S3). Our results also support the hypothesis that the dual feeding regime of *C. s. algoaldensis* is a consequence of its recombinant genome, suggesting the role of hybridization on trophic niche expansion. This ecological novelty could be key to explain the evolutionary success and persistence of *C. s. algoaldensis*, as already documented for other hybrid lineages that have thrived by exploiting novel food resources (Schwarz et al. 2005; Lamichhaney et al. 2018). Although we cannot discard that the dual feeding regime of *C. s. algoaldensis* is the result of post-speciation selection and adaptation, some lines of evidence suggest that this alternative hypothesis is less plausible. Given that all *Chorthippus* species are graminivorous (Gangwere and Morales-Agacino 1973; Gardiner and Hill 2004), it has been suggested a feeding regime based on scrub legumes represents the derived state (Picaud et al. 2002, 2003). Considering that feeding-related traits are likely polygenic, with an underlying complex genetic architecture including genes involved in chemosensory functions, plant recognition, and detoxifying and metabolic pathways (Simon et al. 2015), we argue that the most parsimonious explanation for the dual feeding strategy in *C. s. algoaldensis* is that this trait is an outcome of hybridization rather than the result of a partial evolutionary reversal to the ancestral graminivorous state.

### A biogeographic scenario for Pleistocene hybrid speciation

Pleistocene climatic fluctuations in temperate regions have been hypothesized to increase opportunities for divergence in isolation through distributional shifts and range fragmentation (“species pump” hypothesis; Knowles 2001; *e.g.*, Papadopoulou and Knowles 2015; Ortego and Knowles 2021). Alternatively, it has been proposed that such climatic dynamics could have also prevented speciation through eroding incipient divergences as a result of secondary contact and lineage fusion linked to range expansions during favorable periods (“melting pot” hypothesis; Klicka and Zink 1997; *e.g.*, Maier et al. 2019; Ebdon et al. 2021). According to our divergence time estimates, the clades *C. binotatus* and *C. saulcyi* split during the Gelasian (*ca*. 2.1 Ma; Table 2), which falls within the crown age estimated for the species group *Chorthippus binotatus* based on mtDNA data (*ca*. 1.5-3.0 Ma; García-Navas et al. 2017). As hypothesized for most Gomphocerinae grasshoppers (Mayer et al. 2010; Noguerales et al. 2018b; Ortego et al. 2021), the ancestral diversification of the species complex would be compatible with a scenario of geographical isolation promoted by range fragmentation since the onset of the Pleistocene (Noguerales et al. 2018b; Tonzo et al. 2021). Given the contrasting feeding strategies exhibited by the two main clades, it would be also plausible that adaptive divergence linked to the usage of differing host plants contributed to the early diversification of the group, as documented for other Orthoptera (Apple et al. 2010; Grace et al. 2010). Rather than mutually exclusive, both ecologically-driven divergent selection and allopatric divergence processes could have synergistically contributed to the radiation of *C. binotatus* and *C. saulcyi* clades (Hoskin et al. 2005). Given that the formation of new taxa is generally understood as a protracted process (Rosindell et al. 2010), the time elapsed between the onset of divergence and the identified hybridization event (Calabrian age, *ca*. 1.4 Ma; Table 2) that gave rise to the formation of *C. s. algoaldensis* could be interpreted as an estimate of the pace of speciation in this group (Dynessius and Jansson 2014; Sukumaran et al. 2021). Our coalescent-based estimates indicate that reproductive isolation accumulated since the ancestral split was not enough to prevent interbreeding during at least *ca*. 0.8 Ma (Table 2). This finding would agree with previous studies on insects, for which the minimum time for total hybrid inviability has been estimated in 2-4 Ma (Coyne and Orr 1997; Presgraves 2002). This would support the notion that ancestral diversification likely occurred in geographical isolation, in line with expectations from allopatric speciation models predicting that the completion of reproductive isolation is a more lengthy process when primarily driven by genetic drift (Coyne and Orr 1989).

The magnitude and duration of glacial-interglacial pulses increased during the last stages of the Pleistocene (Head and Gibbard 2005; Hewitt 2011), which led to severe vegetation shifts and repeated phases of reduction in forest extension in favor of grasslands and shrub-like habitats throughout Europe (Donders et al. 2021). Thereby, it is plausible that during this period *C. binotatus* and *C. saulcyi* experienced marked distributional shifts and some of their populations came into secondary contact and hybridized, resulting in the establishment of a hybrid population that rapidly underwent geographic isolation. As expected for hybrid swarms in which population sizes of initial parental demes can be different, the genomic contribution of each parental to the recombinant genome of *C. s. algoaldensis* was uneven (Nolte and Tautz 2010). Our suggested Pleistocene hybrid speciation model would be in concordance with the prediction that temporal changes in habitat distribution and structure increase opportunities of hybridization between formerly isolated lineages (Anderson 1948; Singhal et al. 2021). *In situ* persistence of the nascent hybrid species and its long-lasting geographical isolation from parental lineages could be facilitated by the complex topography characterizing the Central Massif, a region that has been already documented to be an extra-Mediterranean Pleistocene refugia for temperate species (Schmitt and Varga 2012; Kropf et al. 2012; Ursenbacher et al. 2015). Beyond geographic isolation in a topographically complex climate refugium, it is also possible that trophic niche expansion may have conferred an adaptive value to the hybrid lineage, particularly under the highly dynamic environmental conditions prevailing during the Pleistocene that might have resulted in spatial mismatches between the grasshopper climatic niche and host-plant distributions (Noguerales et al. 2018a).

## CONCLUSIONS

We propose a hybrid speciation model where ancient admixture and allopatric isolation in climate refugia can provide a suitable context for hybrid lineages to isolate from parental populations and persist through evolutionary time (James and Abbott 2005; Duenez-Guzman et al. 2009). This two-stage scenario – rapid fusion of parental lineages and isolation of the hybrid swarm – emphasizes the potential importance of Pleistocene-driven demographic dynamics to the formation of homoploid hybrid species. This hypothesis does not neglect the need of evaluating hybridization-derived reproductive isolation (Schumer et al. 2014), but it intends to offer alternative pathways for understanding ancient events of homoploid hybrid speciation in which demonstrating that hybridization triggered reproductive isolation with parental lineages is challenging or virtually impossible (Nieto-Feliner et al. 2017; Edelman and Mallet 2021). Collectively, our study provides insights on how the interplay between extended speciation duration and climate-mediated increasing opportunities for gene flow may eventually promote hybrid speciation (Dynessius and Jansson 2014). As opposed to the classic view of melting pots preventing diversification (Klicka and Zink 1997; Ebdon et al. 2021), our results offer more nuanced insights on Pleistocene speciation by highlighting how a combination of allopatric divergence and subsequent hybridization can both contribute to diversification (Seehausen et al. 2014; Meier et al. 2017; Gillespie et al. 2020).

## AUTHORS CONTRIBUTIONS

VN and JO conceived and designed the study and analyses. VN and JO collected the samples. VN analyzed the data and wrote the manuscript with inputs from JO.

## ACKNOWLEDGMENTS

We are grateful to Pedro J. Cordero and Amparo Hidalgo-Galiana for their help during field and laboratory work, respectively. We wish to thank to Centro de Supercomputación de Galicia (CESGA) and Doñana’s Singular Scientific-Technical Infrastructure (ICTS-RBD) for access to computer resources. Logistical support was provided by Laboratorio de Ecología Molecular (LEM-EBD) from Estación Biológica de Doñana. This work was funded by the Spanish Ministry of Economy and Competitiveness and the European Regional Development Fund (grants CGL2011-25053, CGL2014-54671-P, CGL2016-80742-R, and CGL2017-83433-P). During this work, VN was supported by a postdoctoral contract under the iBioGen project (grant agreement No 810729), which has received funding from the European Union’s Horizon 2020 research and innovation programme, and by a Juan de la Cierva-Formación postdoctoral fellowship (grant FJC2018-035611-I) funded by MCIN/AEI/10.13039/501100011033.

## DATA ARCHIVING

Raw Illumina reads will be deposited at the NCBI Sequence Read Archive (SRA) upon manuscript acceptance. Input files for all analyses will be available for download from Figshare upon manuscript acceptance.

## SUPPORTING INFORMATION

**Table S1.**
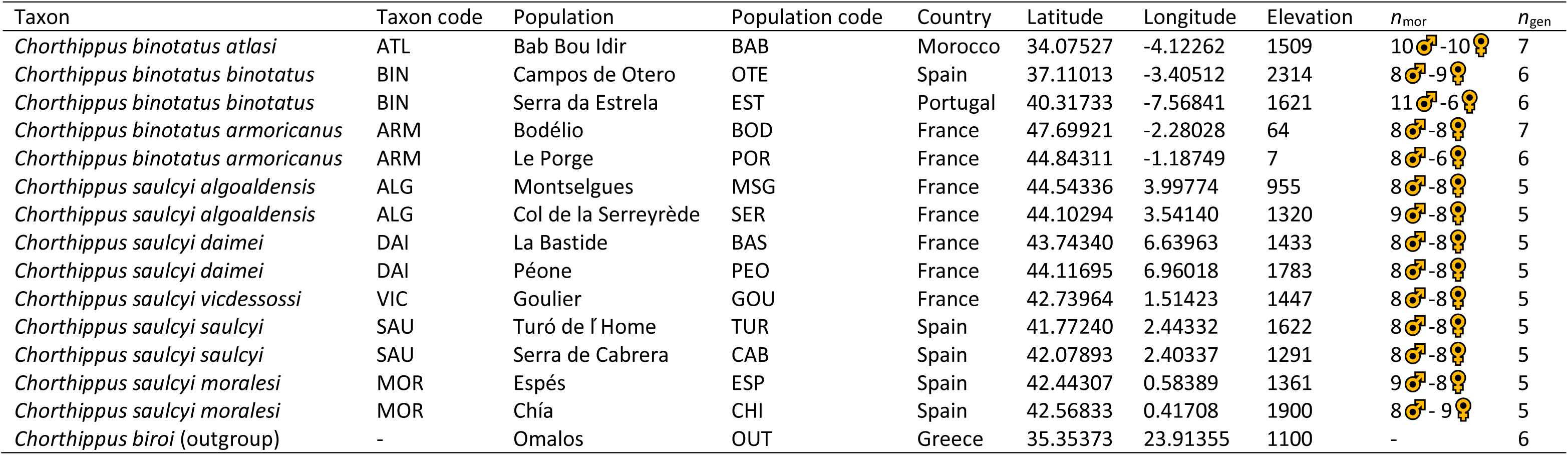
Geographical location and taxonomic information of the sampled populations from the studied species complex. The number of analyzed individuals per population for genetic (n_gen_) and morphological (n_mor_) analyses are indicated.

**Table S2.** Analyses of introgression using *D*-statistic (ABBA/BABA) tests performed considering all possible subspecies combinations for *C. binotatus* (P_3_) and *C. saulcyi* (P_1_). The standard deviation (SD) of the *D*-statistic (*D*) was estimated for each comparison with 1000 bootstrap replicates. The number of informative sites (*n* loci) used in each comparison is also shown. *Chorthippus biroi* was used as an outgroup taxon. Taxon codes as in Table S1.

| $P_1$ | $P_2$ | $P_3$ | BABA sites | ABBA sites | $D(\pm \text{SD})$ | Z-score | P-value | *n* loci |
| --- | --- | --- | --- | --- | --- | --- | --- | --- |
| SAU | ALG | ATL | 43 | 85 | $0.33 \pm 0.10$ | 3.36 | <0.001 | 525 |
| MOR | ALG | ATL | 46 | 82 | $0.28 \pm 0.11$ | 2.53 | 0.011 | 541 |
| VIC | ALG | ATL | 44 | 67 | $0.20 \pm 0.14$ | 1.40 | 0.161 | 471 |
| DAI | ALG | ATL | 45 | 85 | $0.31 \pm 0.12$ | 2.61 | 0.009 | 539 |
| SAU | ALG | BIN | 59 | 84 | $0.18 \pm 0.09$ | 1.97 | 0.048 | 1076 |
| MOR | ALG | BIN | 50 | 86 | $0.26 \pm 0.08$ | 3.28 | <0.001 | 1062 |
| VIC | ALG | BIN | 36 | 60 | $0.26 \pm 0.10$ | 2.65 | 0.008 | 840 |
| DAI | ALG | BIN | 40 | 94 | $0.40 \pm 0.08$ | 4.83 | <0.001 | 1023 |
| SAU | ALG | ARM | 48 | 127 | $0.45 \pm 0.07$ | 6.28 | <0.001 | 1041 |
| MOR | ALG | ARM | 48 | 123 | $0.44 \pm 0.08$ | 5.85 | <0.001 | 1025 |
| VIC | ALG | ARM | 41 | 123 | $0.50 \pm 0.08$ | 6.33 | <0.001 | 988 |
| DAI | ALG | ARM | 34 | 83 | $0.41 \pm 0.09$ | 4.44 | <0.001 | 816 |

**Table S3.** Results of HYDE analyses testing for hybridization in C. s. algoaldensis considering all possible subspecies combinations for parental taxa C. saulcyi (P_1_) and C. binotatus (P_3_). The putative hybrid lineage (P_HYB_) is modeled as a mixture between two parental populations, P_1_ (1-γ) and P_3_ (γ). Confidence interval (95 % CI) of the γ parameter was estimated from 100 bootstrap resampling of the individuals within the putative hybrid lineage. We also reported the minimum and maximum values (range) of the γ parameter across putatively hybrid individuals (γ_IND_) according to the results of individual-based analyses. Analyses were run using 16,220 sequence loci, with a total of ∼2.02 M bp length. Chorthippus biroi was used as an outgroup taxon. Taxon codes as in Table S1.

| $P_1$ | $P_{HYB}$ | $P_3$ | $\gamma$ (95% CI) | Z-score | P-value | $\gamma_{IND}$ range |
| --- | --- | --- | --- | --- | --- | --- |
| SAU | ALG | ATL | 0.33 (0.32-0.34) | 73.75 | <0.001 | 0.30-0.35 |
| MOR | ALG | ATL | 0.59 (0.58-0.61) | 47.69 | <0.001 | 0.56-0.75 |
| VIC | ALG | ATL | 0.60 (0.58-0.65) | 33.15 | <0.001 | 0.56-0.68 |
| DAI | ALG | ATL | 0.51 (0.50-0.52) | 41.00 | <0.001 | 0.48-0.63 |
| SAU | ALG | BIN | 0.32 (0.31-0.34) | 65.89 | <0.001 | 0.28-0.36 |
| MOR | ALG | BIN | 0.62 (0.60-0.64) | 44.15 | <0.001 | 0.58-0.74 |
| VIC | ALG | BIN | 0.61 (0.59-0.64) | 40.36 | <0.001 | 0.57-0.78 |
| DAI | ALG | BIN | 0.53 (0.52-0.54) | 46.49 | <0.001 | 0.51-0.59 |
| SAU | ALG | ARM | 0.31 (0.29-0.33) | 51.64 | <0.001 | 0.24-0.35 |
| MOR | ALG | ARM | 0.68 (0.65-0.72) | 34.38 | <0.001 | 0.63-0.91 |
| VIC | ALG | ARM | 0.64 (0.62-0.66) | 42.65 | <0.001 | 0.60-0.78 |
| DAI | ALG | ARM | 0.57 (0.56-0.58) | 48.87 | <0.001 | 0.54-0.64 |

**Table S4.** Results of the analyses testing for differences in (i) forewing length relative to femur length (FWL/FL), (ii) forewing median area length relative to total forewing length (MAL/FWL), and (iii) prozone length relative to total pronotum length (PZ/PR) between the focal taxon *Chorthippus saulcyi algoaldensis* and *C. binotatus* and *C. saulcyi*, using post-hoc Tukey ’s tests. Adjusted *P*-values are reported, presented below the diagonal for males and above the diagonal for females, with values in bold indicating significant comparisons.

| FWL/FL |  |  |  |  |
| --- | --- | --- | --- | --- |
|  |  | *C. binotatus* | *C. s. algoaldensis* | *C. saulcyi* |
|  | *C. binotatus* | - | **<0.001** | **<0.001** |
|  | *C. s. algoaldensis* | **<0.001** | - | 0.789 |
|  | *C. saulcyi* | **<0.001** | 0.750 | - |
| MAL/FWL |  |  |  |  |
|  |  | *C. binotatus* | *C. s. algoaldensis* | *C. saulcyi* |
|  | *C. binotatus* | - | **<0.001** | **<0.001** |
|  | *C. s. algoaldensis* | **<0.001** | - | **0.025** |
|  | *C. saulcyi* | **<0.001** | 0.290 | - |
| PZ/PR |  |  |  |  |
|  |  | *C. binotatus* | *C. s. algoaldensis* | *C. saulcyi* |
|  | *C. binotatus* | - | **0.014** | **<0.001** |
|  | *C. s. algoaldensis* | **<0.001** | - | 0.400 |
|  | *C. saulcyi* | **<0.001** | 0.753 | - |

**Table S5.**
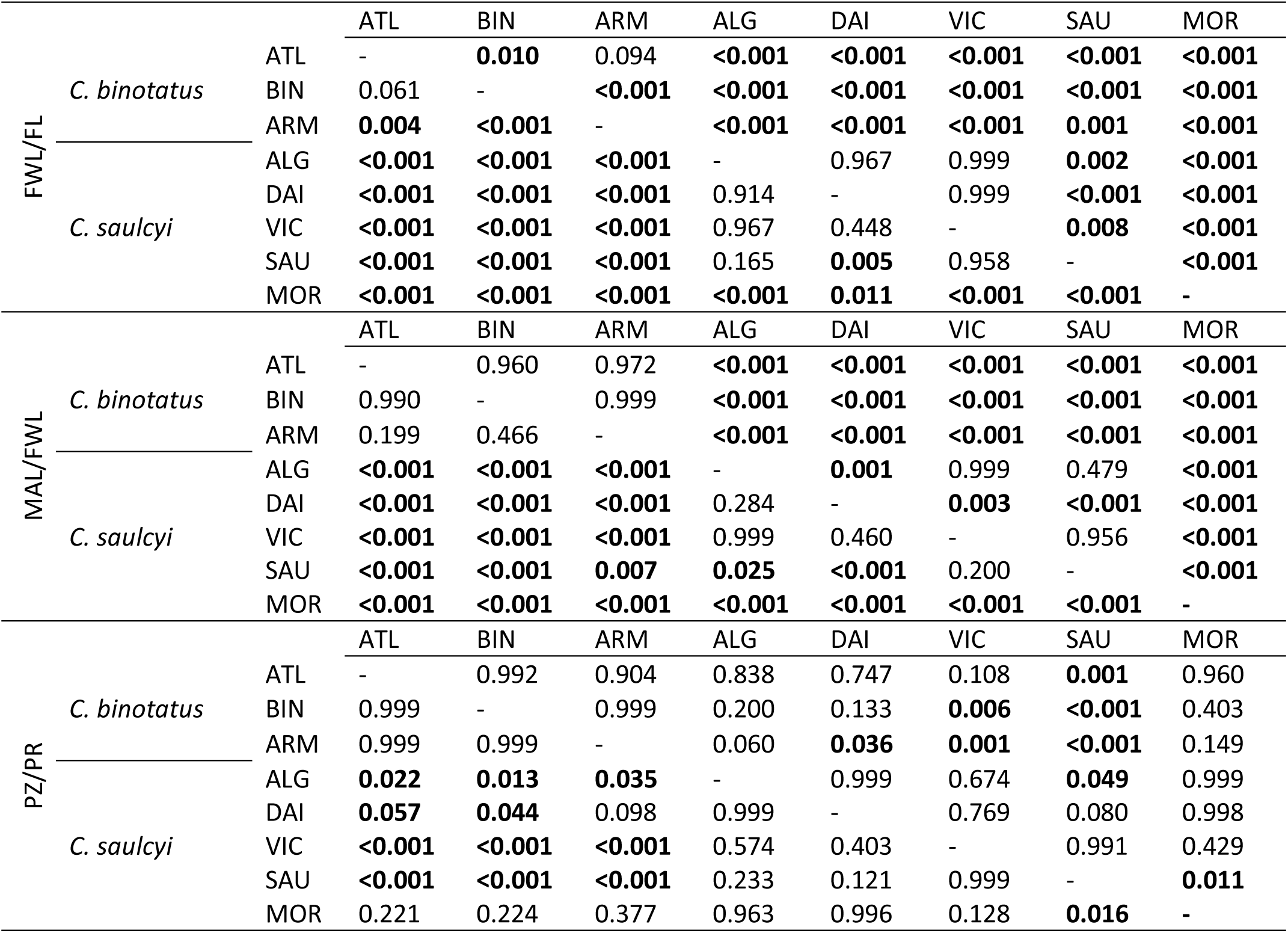
Results of the analyses testing for differences in (i) forewing length relative to femur length (FWL/FL), (ii) forewing median area length relative to total forewing length (MAL/FWL), and (iii) prozone length relative to total pronotum length (PZ/PR) between the different subspecies of the studied species complex, using post-hoc Tukey ’s tests. Adjusted P-values are reported, presented below the diagonal for males and above the diagonal for females, with values in bold indicating significant comparisons. Subspecies codes as in Table S1.

**Table S6.** Mahalanobis distances (*D*) between the focal taxon *Chorthippus saulcyi algoaldensis* and *C. binotatus* and *C. saulcyi* obtained through a Canonical Variates Analysis (CVA) for forewing shape. Values for males are presented below the diagonal and for females above the diagonal. Values in bold indicate significant Mahalanobis distances (*D*).

|  | *C. binotatus* | *C. s. algoaldensis* | *C. saulcyi* |
| --- | --- | --- | --- |
| *C. binotatus* | - | **4.372** | **4.557** |
| *C. s. algoaldensis* | **5.412** | - | **2.031** |
| *C. saulcyi* | **6.252** | **2.972** | - |

**Table S7.** Mahalanobis distances (D) between the different subspecies of the studied species complex obtained through a Canonical Variates Analysis (CVA) for forewing shape. Values for males are presented below the diagonal and for females above the diagonal. Values in bold indicate significant Mahalanobis distances (D). Subspecies codes as in Table S1.

|  |  | ATL | BIN | ARM | ALG | DAI | VIC | SAU | MOR |
| --- | --- | --- | --- | --- | --- | --- | --- | --- | --- |
| *C. binotatus* | ATL | - | **3.813** | **4.358** | **6.431** | **7.866** | **5.825** | **5.133** | **9.841** |
|  | BIN | **2.538** | - | **2.616** | **6.028** | **7.277** | **5.480** | **4.859** | **9.004** |
|  | ARM | **2.521** | **2.227** | - | **5.148** | **6.149** | **4.580** | **4.401** | **7.994** |
| *C. saulcyi* | ALG | **7.880** | **6.716** | **7.072** | - | **2.886** | **3.708** | **3.034** | **4.431** |
|  | DAI | **9.276** | **8.302** | **8.337** | **3.201** | - | **4.360** | **4.330** | **3.350** |
|  | VIC | **7.382** | **6.700** | **6.849** | **3.956** | **3.110** | - | **2.580** | **6.060** |
|  | SAU | **6.754** | **6.103** | **6.207** | **3.592** | **3.658** | 1.807 | - | **6.258** |
|  | MOR | **11.796** | **10.587** | **10.665** | **5.318** | **3.974** | **6.054** | **6.331** | - |

**Figure S1.**
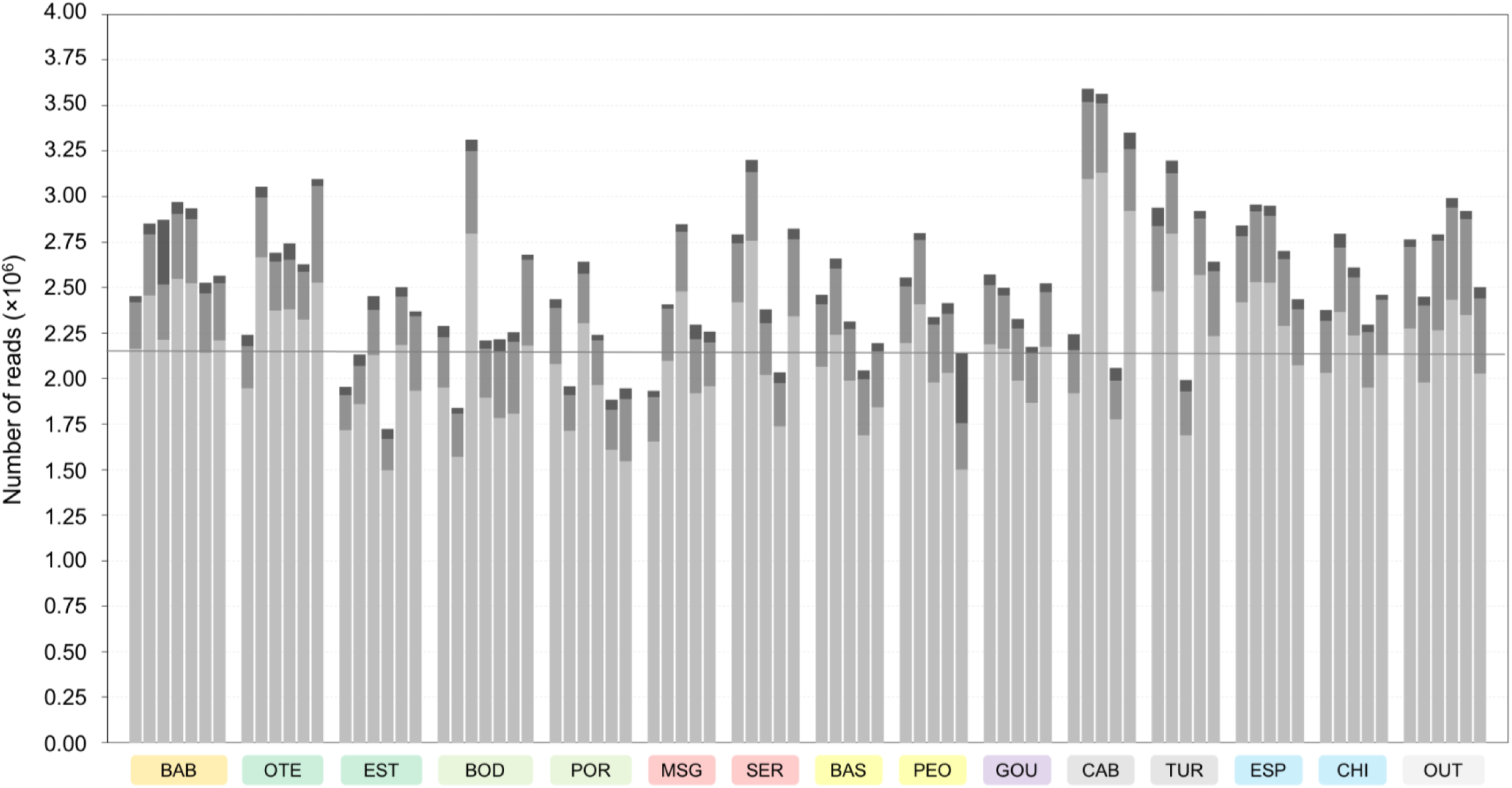
Number of reads per individual before and after different quality filtering steps by STACKS and PYRAD. The cumulative stacked bars represent the total number of raw reads obtained for each individual. Within each bar, the dark grey color represents the reads that were discarded by STACKS (*process_radtags*) due to low quality, adapter contamination or an ambiguous barcode. Medium grey color represents the reads that were subsequently discarded by PYRAD after filtering out reads that did not comply with the quality criteria (reads with >2 sites with a Phred quality score <20 were discarded). Finally, light grey color represents the total number of retained reads used to identify homologous loci during the subsequent steps performed in PYRAD. Grey horizontal line indicates the average number of reads across all individuals (*n* = 83). Population codes as in Table S1.

**Figure S2.**
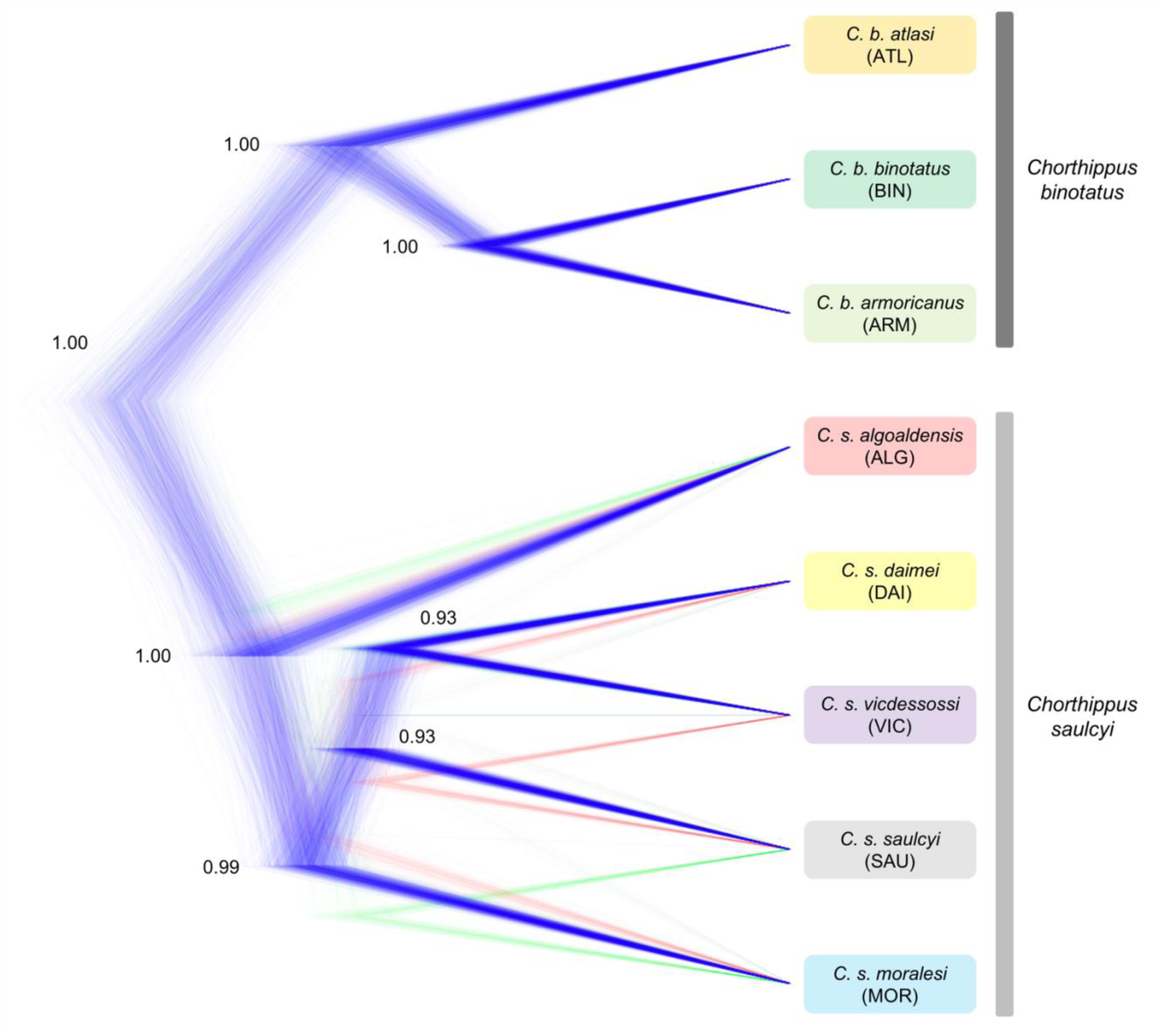
Species tree as inferred by SNAPP (based on 2,926 SNPs) displaying the phylogenetic relationships among taxa from the studied species complex. Posterior probabilities for the most supported topology are indicated on the nodes. Taxon codes as in Table S1.

**Figure S3.**
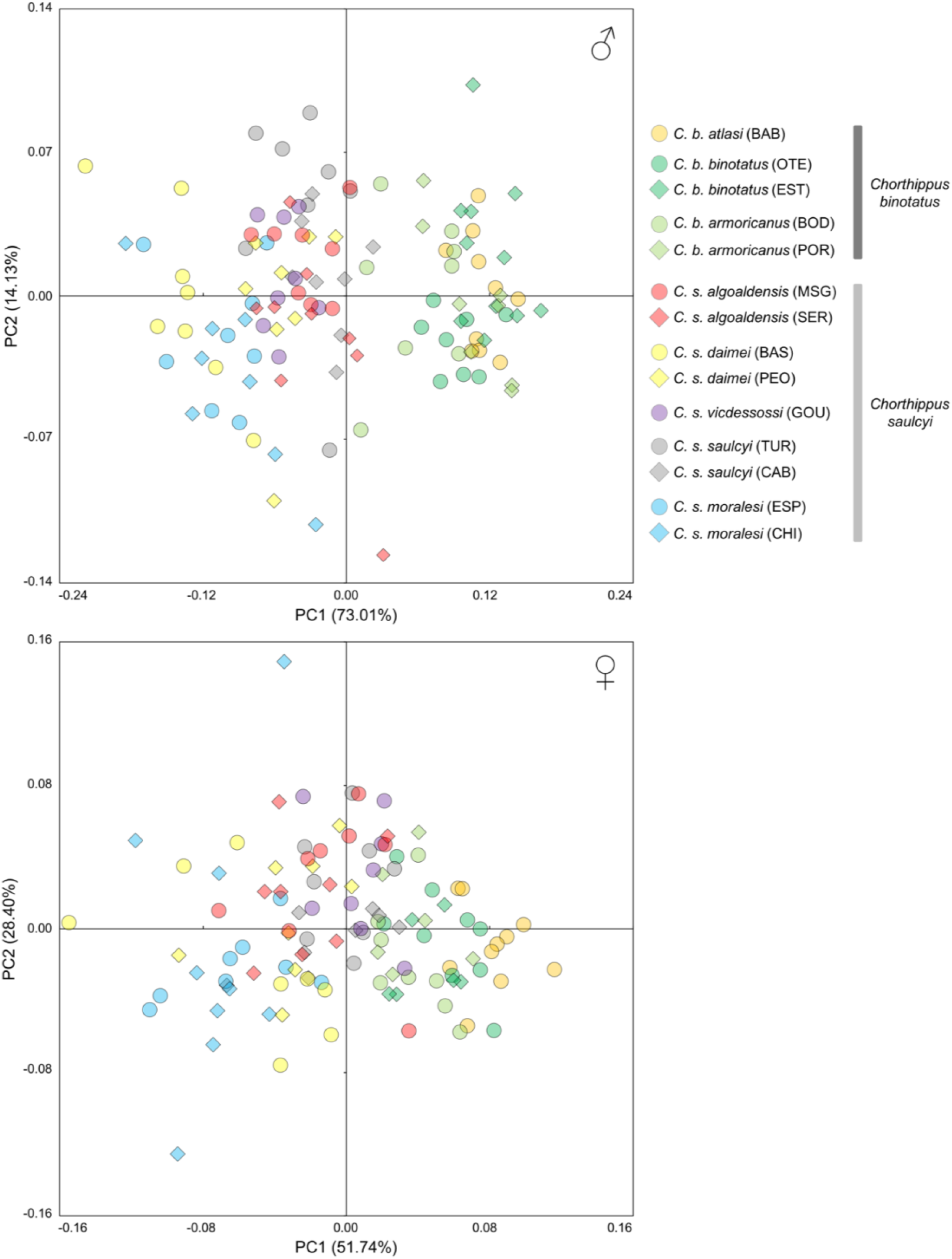
Forewing shape variation in males (top panel) and females (bottom panel) for the different subspecies and populations of the studied complex along the two first principal components (PCs). Population codes as in Table S1.

### Methods S1. Genomic library preparation

We used a salt extraction protocol to purify genomic DNA from a hind femur of each specimen (Aljanabi and Martinez 1997). Genomic DNA from each specimen was individually barcoded and processed into a genomic library using the double digestion restriction associated DNA sequencing (ddRADseq) procedure described in Peterson et al (2012) with some minor modifications as detailed in Lanier et al. (2015) and Massatti and Knowles (2016). Briefly, DNA was double-digested using *Eco*RI and *Mse*I restriction enzymes (New England Biolabs), followed by the ligation of Illumina adaptors and unique 7-base-pair barcodes. Ligation products were pooled, size-selected between 475 and 580 base pairs (bp) using a Pippin Prep (Sage Science) machine, and amplified by iProof^TM^ High-Fidelity DNA Polymerase (BIO-RAD) with 12 cycles. Single-read 151 bp sequencing was performed on an Illumina HiSeq2500 platform at The Centre for Applied Genomics (Hospital for Sick Children, Toronto, Canada).

### Methods S2. Genomic data filtering and sequence assembly

Raw sequence reads were demultiplexed and quality-filtered using the *process_radtags* script within the STACKS v.1.35 pipeline (Catchen et al. 2013). Only reads with a Phred score >10 (using a sliding window of 15%), no adaptor contamination, and that had an unambiguous barcode and restriction cut site were retained. Then, we checked the quality of raw sequences in FASTQC v.0.11.5 (http://www.bioinformatics.babraham.ac.uk/projects/fastqc/) and trimmed them to 129 bp using SEQTK (Heng Li, https://github.com/lh3/seqtk) in order to remove low-quality reads near the 3′ ends. Reads retained after *process_radtags* were further quality-filtered using the program PYRAD v.3.0.66 (Eaton 2014) to convert base calls with a Phred score <20 into Ns and discard reads with >2 Ns (Fig. S1). Afterwards, we used PYRAD to cluster retained reads within- and across samples considering a clustering threshold of sequence similarity of 85% (*W*_CLUST_ = 0.85). Clusters with a minimum coverage depth less than 5 (*d* = 5) and loci containing one or more heterozygous sites across more than 15% of individuals (*maxSH* = p.15) and more than 20 polymorphic sites (*maxSNPs* = 20) were discarded (Eaton 2014; Hipp et al. 2014). In a final filtering step we retained those loci that were present in at least 25% of the samples (minimum taxon coverage, *minCov* = 25%; Noguerales et al. 2018), which retained a total of 16,119 unlinked SNPs.

